# Resolving near-micron scale features within a whole sheep head using Hierarchical Phase-Contrast Tomography

**DOI:** 10.64898/2025.12.15.694379

**Authors:** Theresa Urban, Joseph Brunet, Marius Didziokas, Matthieu Chourrout, Andrew Keenlyside, Karla Orsine Murta Dias, Alessandro Mirone, Alexandre Bellier, Hector Dejea, Laurent Lammalle, Bernadette de Bakker, Harry Petrushkin, Mehran Moazen, Claire L. Walsh, Peter D. Lee, Paul Tafforeau

## Abstract

**Background:** Hierarchical Phase-Contrast Tomography (HiP-CT) was developed to image *ex vivo* intact human soft-tissue organs with local near-micron scale resolution.

**Purpose:** We demonstrate the application of HiP-CT in combination with the recently developed Eikonal Phase Retrieval (EPR) for resolving anatomical features in large hard and soft tissue structures on a sheep head, and show the applicability for zooming to resolve anatomical features with near-micron scale resolution.

**Materials and Methods:** We imaged an entire skinned sheep head prepared in 70 % ethanol with HiP-CT at an isotropic voxel size of 16.5 µm using the recently developed Eikonal Phase Retrieval (EPR) to reduce artefacts from bones. Local tomography zooms were taken in regions of interest: the eye (4.23 µm voxels) and the coronal suture (2.20 µm voxels). To compare to clinical imaging, we acquired T2-weighted MRI and CT of the sheep head and evaluated the contrast-to-noise ratio for differentiation of white and grey matter in the brain for all modalities. Further evaluations on HiP-CT images include structure tensor analysis for structural orientation in the brain as well as fibre tracking in the coronal suture.

**Results:** HiP-CT combined with EPR achieved high contrast for both hard and soft tissue. Comparison to MRI showed similar soft-tissue contrast, but much higher spatial resolution. Structure tensor analysis in the brain revealed the orientation of the major white matter bundles. In the eye, near-micron scale features such as retinal layers and the bundles in the optic nerve were visualized. Fiber tracking allowed analysis of the orientation of collagen fiber bundles in the coronal suture.

**Conclusion:** This work highlights the potential of HiP-CT to image a complete sheep head, ex vivo, and hierarchically zoom without sectioning to resolve few-microns features locally, enabling comprehensive three-dimensional visualization of intricate cranial structures and their spatial interrelations.

**Summary statement:** Technical developments in HiP-CT enable ex vivo X-ray imaging of an intact sheep head with local resolution to near-micron scale, demonstrating future viability on a human head.

**Key results:**

1. A whole sheep head was imaged with synchrotron-based hierarchical phase-contrast tomography coupled with Eikonal Phase-retrieval with 16.5 µm isotropic voxels.
2. The high contrast for both hard and soft tissue enabled differentiation of brain white and grey matter, the optic nerve and bone features within the skull.
3. Local tomography zoom scans in an eye (4.23 µm voxels) and coronal suture (2.20 µm voxels) allowed visualization and analysis of near-micron sized features.

## Introduction

The head is one of the most complex regions of the body, where the brain, sensory and endocrine organs, vessels, and other vital structures are densely packed into a confined space. Understanding these structures’ intricate three-dimensional microanatomy and spatial relationships is crucial for a wide range of researchers and clinicians.

A range of imaging techniques serve this purpose, offering differing contrast modalities, fields of view, resolutions, and in-vivo compatibility, but all with limitations. Clinical methods, including magnetic resonance imaging (MRI) and computed tomography (CT), offer a whole-head coverage *in vivo*, but have limited resolution (hundreds of microns)^1,2^. *Ex vivo* methods, e.g. micro-CT (µCT) offer high resolution (around 1 µm)^3^ but sample size is usually limited to a few centimeters and often requires staining for soft tissue contrast. Sectioning methods like histology or volume electron microscopy^4^ allow single-cell and higher resolutions but are destructive and again usually limited to a few centimeters sample size.

Synchrotrons are large-scale facilities that provide X-rays suitable for high resolution and high contrast imaging of a wide range of biological and biomedical ex-vivo samples. Synchrotron-based Hierarchical Phase-Contrast Tomography (HiP-CT)^5^ was recently developed for imaging intact human organs. HiP-CT involves an initial overview scan of the complete sample (∼15 cm diameter for most organs) with typical voxel sizes of ca. 20 µm, followed by local tomography zooms targeting volumes of interest of a few cubic cm with voxel sizes down to ca. 2 µm, without biopsy. The technique has been applied to various organs^6–9^ within the Human Organ Atlas project^10^.

Previous studies with HiP-CT on large samples were all only soft tissue. Imaging large heterogeneous samples containing both soft and hard tissue, such as the head, poses challenges on both image acquisition and reconstruction. Conventional phase retrieval methods, like Paganin’s algorithm^11^, make simplifying assumptions that are not valid in this case, introducing streak artefacts and halos. To overcome these limitations, an iterative method for phase retrieval, Eikonal Phase Retrieval (EPR), was recently introduced^12^.

In this work, we couple HiP-CT and EPR, presenting the technical developments required to image an entire sheep head (16.5 µm voxels) and zoom locally to demonstrate how it can reveal previously unseen anatomical features with near-micron scale resolution and high contrast (e.g. 2.20 µm voxel tracking of collagen fiber bundles within the coronal suture).

## Methods

### Sample

The sheep head was bought already skinned at a local butchery. It was prepared according to the standardized protocol for HiP-CT^13^. After infusion of an embalming fluid (GENELYN Artériel Ultra, EEP Company, France) via the carotid arteries and subsequent fixation in 4% formalin it was transferred to 70 % ethanol via a series of ethanol baths (50 %, 60 %, 70 %, 70 %). It was then vacuum degassed and mounted in a 15 cm diameter sample container with agar gel support. The sample container was further degassed with an inline-degassing system before imaging.

### HiP-CT Imaging – Overview Scans

Two overview scans with HiP-CT were acquired at ESRF-EBS beamline BM18, as introduced by Walsh et al.^5^. Beam parameters and processing were adapted to large samples with mixed soft and hard tissues (see Fig. 1 and Suppl. Table 1). The overall imaged volume has a diameter of 14.9 cm and a height of 22 cm.

**Figure 1:**
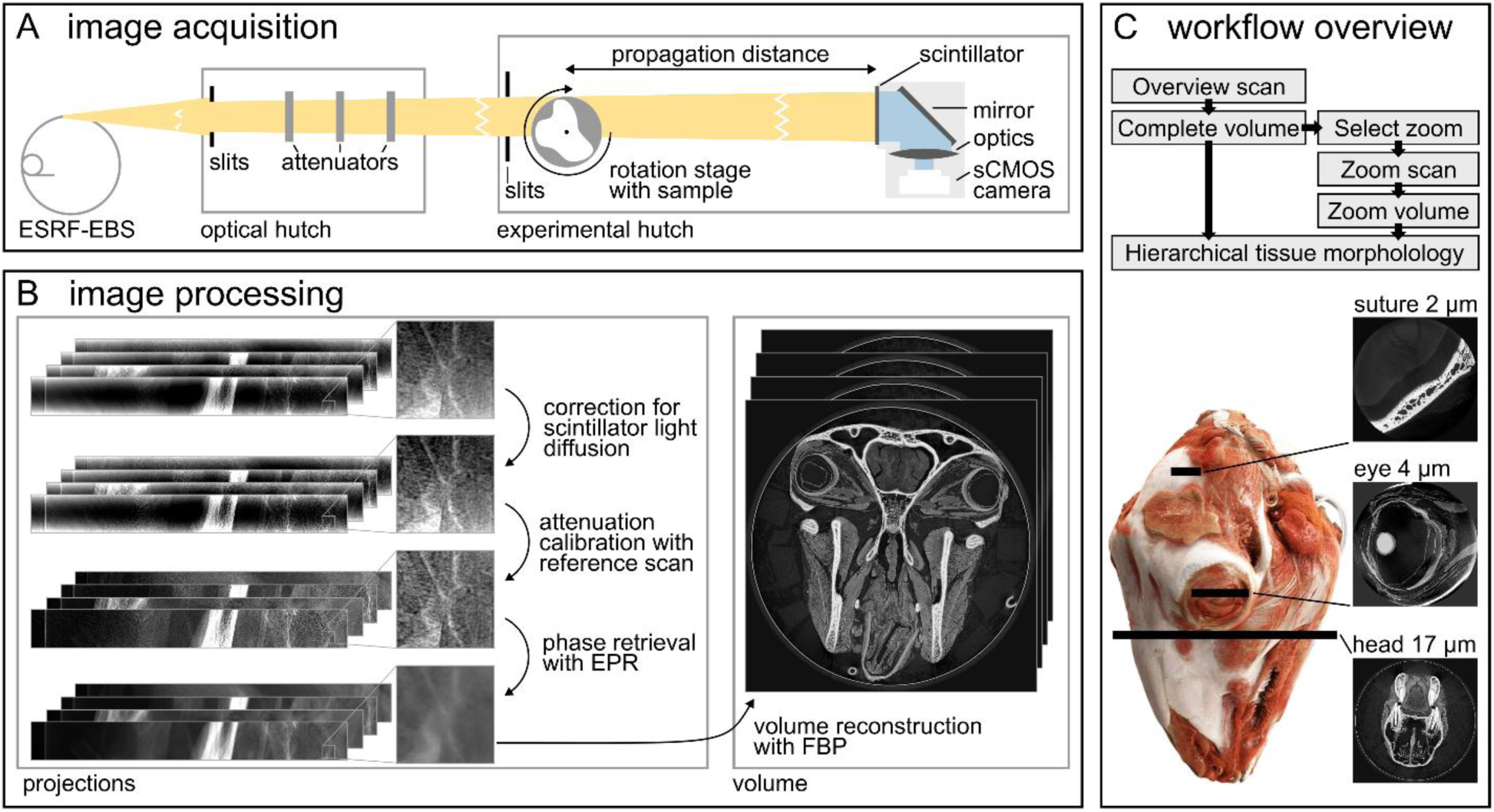
HiP-CT imaging. A, schematic view of the ESRF-EBS beamline BM18 with its main components. Besides slits and attenuators, there are no optical components in the beam path. The propagation distance is adapted to the pixel size of the image acquisition and ranges between 1.5 m (zoom with 2.2 µm voxels) and 30 m (overview scan with 16.5 µm voxels). A different scintillator and optic are used for overview and zoom scan. B, image reconstruction pipeline for the overview scan. All projections are corrected for scintillator light diffusion, and calibrated with the attenuation-matched reference scan before phase retrieval with EPR and volume reconstruction. C, overall workflow. Zoom volumes are selected from the overview scan, enabling hierarchical tissue morphology analysis.

To reduce beam hardening and improve detector dynamic all projections were calibrated against a reference scan acquired from an identical container filled only with 70 % ethanol^13^. Phase retrieval was performed on the projections with the iterative EPR algorithm. EPR accounts for a polychromatic X-ray spectrum, uses a forward propagation from sample to detector, and includes a deconvolution step to correct for visible light diffusion within the scintillator, reducing artefacts from bones^12^. Volume reconstruction was performed with the inhouse reconstruction pipeline night_rail^14^, using a filtered backprojection algorithm implemented in Nabu^15^, followed by ring artefact correction^16^.

### HiP-CT Imaging – Local Tomography Zoom Scans

Regions of interest were selected from the overview scan to perform local tomography zooms at higher resolution without the need for physical sectioning. A total of twelve regions were imaged (see Suppl. Table 1). Two zoom scans are analysed in this study: one capturing the complete eye (voxel size 4.23 µm; volume diameter 38 mm, height 41 mm), and one in the coronal suture (voxel size 2.20 µm; volume diameter 20 mm, height 22 mm). To reduce local tomography artefacts, the references were again taken in a matching ethanol-filled reference container, following the HiP-CT protocol for off-axis zoom scans^5^. Volume reconstruction was performed with the inhouse reconstruction pipeline night_rail^14^. Since the effect of the heterogeneous sample is less pronounced with higher resolutions, Nabu’s^15^ implementation of Paganin’s algorithm^11^ coupled to 2D unsharp mask, was used for phase retrieval.

### Clinical Imaging

To compare HiP-CT with conventional clinical imaging, *ex vivo* MRI and CT scans were also acquired. MRI was performed on a 3 T Achieva dStream (Philips Medical Systems, The Netherlands) 14 months before the 16.5 µm overview scan, using a T2-weighted contrast and a brain imaging sequence with an echo train length of 60 and overall scan duration of 86 min. The reconstructed voxel size was 214×214×225 µm^3^.

CT imaging was performed on an Aquilion ONE (Canon Medical Systems, Japan) machine 2 months after HiP-CT imaging using a head image protocol. In the reconstruction, a convolution kernel for bone was used. The reconstructed voxel size was 460×460×800 µm^3^.

## Evaluations

### 3D Renderings

For rendering of the overview scan, the whole head was segmented from the background using Organ-masker^17^. HiP-CT datasets were binned and rendered using commercial software (Cinematic Anatomy, Siemens Healthineers, Germany) optimized for medical image rendering.

### Brain Analysis

The brain was manually segmented from the complete head with 3D Slicer 5.8.1. The structure tensor over the whole volume of the brain was computed using a width of σ_ST_ = 1 pixel with Cardiotensor^18^. The vector field was reoriented with respect to the anterior commissure—posterior commissure landmarks^19^, as used in human neuroscience.

### CNR Calculation

For comparison of modalities, both MR and CT images were registered to the HiP-CT scan with an affine transformation using Python and SimpleITK^20^. For evaluation of the soft-tissue contrast in the brain, the CNR was computed for all three imaging modalities using identical regions, transferred between modalities with the transforms from the registration. See Suppl. Materials for more details and Suppl. Figure S1 for the transferred segmentations.

### Coronal Suture Analysis

The analysis of the coronal suture was performed using Avizo 2023.1 (Thermo Fisher Scientific, USA). Cranial bone and right coronal suture were segmented from the 2.20 µm local tomography scan^21^, and the thickness of the suture was measured using the thickness map module^22^.

For fibre bundle analysis, three sub-volumes (200×200×200 voxels at 2.20 µm) were extracted from the suture at the dorsal, middle and ventral suture locations of the lateral middle of the coronal suture. The trace correlation line module was used to obtain individual fibre bundles. To confirm the efficacy of the proposed fibre bundle tracing methodology, the periosteum (400×400×400 voxels at 2.20 µm) at the same lateral position was also analysed with the same parameters. Details on segmentation and fibre bundle analysis can be found in the Supplementary Materials.

## Results

Over the course of several months, we acquired two overview HiP-CT scans with 27.7 µm and 16.5 µm voxel size, respectively, various zoom scans with voxel sizes between 4.2 µm and 2.2 µm, a clinical MR and a clinical CT scan of the head. An overview of all HiP-CT scans with their respective context and acquisition parameters is provided in Suppl. Table 1. All imaging data is made publicly available on BioImage Archive.

### Whole-head imaging

An overview of the data from the complete head scan with a voxel size of 16.5 µm is shown in Figure 2, and a video rendering the whole head is available in the Supplementary Material. Imaging the entire intact head enables simultaneous visualization of both the context anatomical features and their spatial interconnections across the whole sample. Both hard and soft tissues were clearly visualized without sectioning. The high soft-tissue contrast provided by phase-contrast imaging coupled with the EPR enables the visualization of features such as the fibers in the tongue (Fig. 2I), the muscles in the jaw (Fig. 2G), and the ciliary body of the eye (Fig. 2J). Notably, white and gray matter in the brain can be clearly differentiated despite the brain being enclosed within the skull. The bone itself, including cranial sutures, can be clearly distinguished with very high contrast and spatial resolution (Fig. 2B, H).

**Figure 2:**
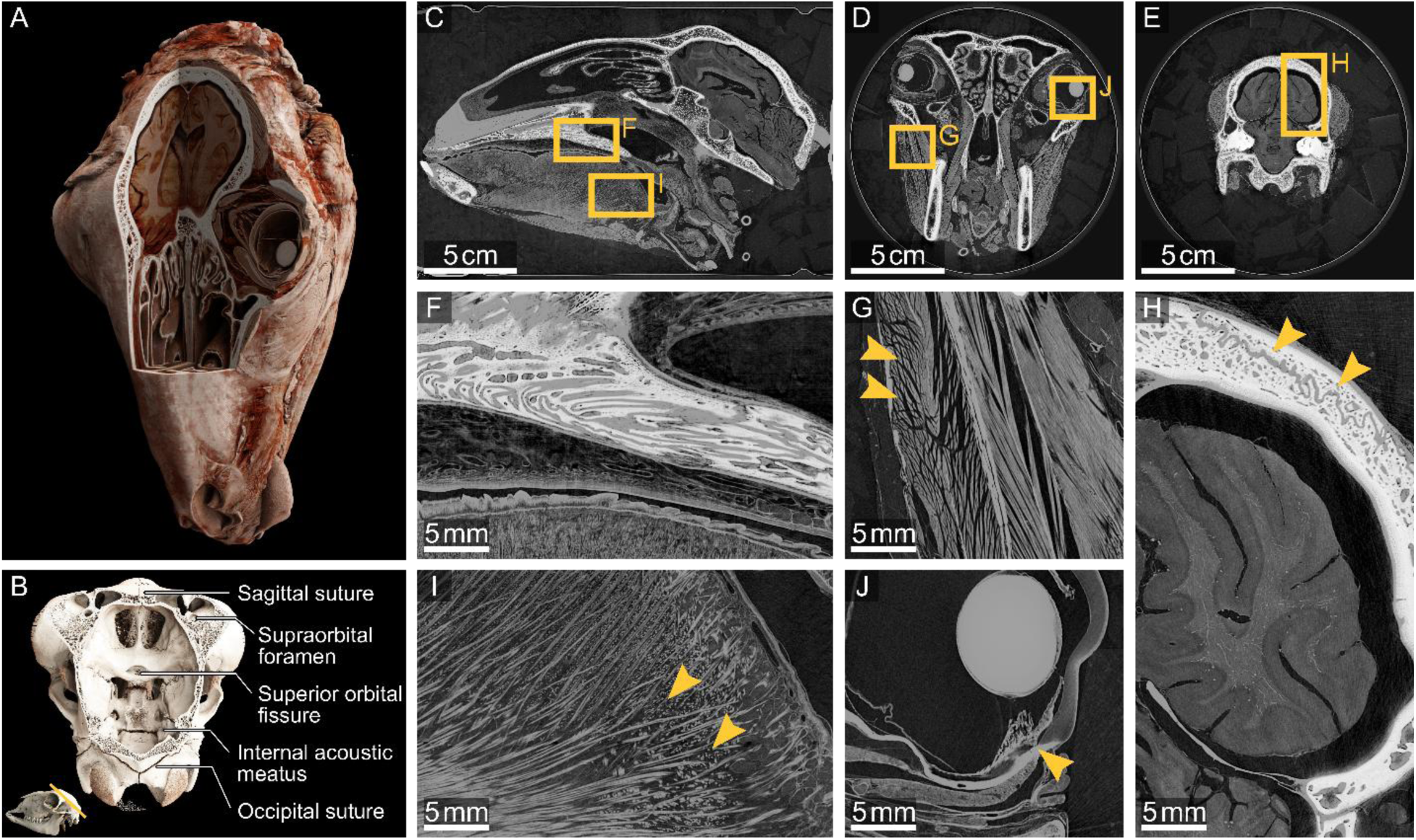
Data overview of the complete sheep’s head with a voxel size of 16.5 µm. A, Volume rendered in Cinematic Anatomy. B, Rendering and annotations of the virtually opened superior view of the skull base. C-J, detailed views to the regions indicated by the rectangles. It is possible to visualize both hard tissue (e.g. palatinum (F), sutures in the skull (H)) and soft tissue (e.g. muscles in the jaw (G) and tongue (I), eye (J), brain (H)).

### Comparison to clinical methods

Figure 3 shows a comparison of the HiP-CT overview scan (16.5 µm voxels) with clinical MRI and CT scans. A video comparing HiP-CT to MR is available in the Supplementary Material.

**Figure 3:**
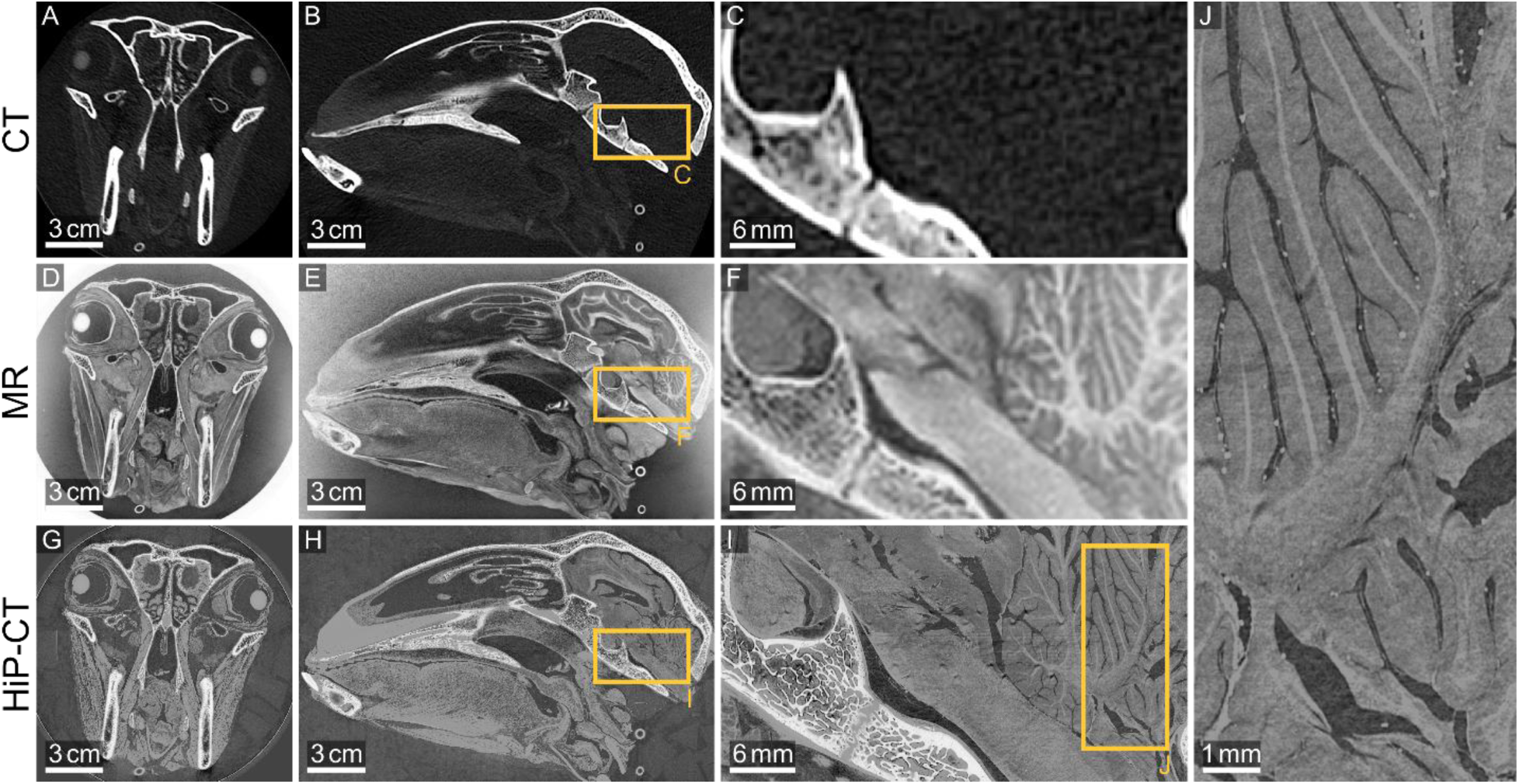
Comparison of HiP-CT with established whole-head clinical imaging methods. A-C, axial and sagittal slices and detail view of the CT scan. D-F, axial and sagittal slices and detail view of the MRI scan in negative contrast for better visual comparison. G-J, Axial and sagittal slices and detail view of the HiP-CT scan at 16.5 µm voxel size. The HiP-CT scan offers high contrast on both hard and soft tissue, and a much higher resolution than the clinical modalities.

HiP-CT with ethanol preparation yields soft-tissue contrast rivalling that of MR (Fig. 3F,I), yet with a much greater spatial resolution. This allows for detailed description of mesoscale structures within the cerebellum (Fig. 3J). HiP-CT more clearly identifies spaces of cerebrospinal fluid around the brainstem while maintaining the capacity to separate isodense soft tissue in the anterior and posterior pituitary gland (Fig. 3I). Similarly, HiP-CT provides high contrast in bones similar to CT but again with much higher spatial resolution, demonstrating the complex internal structure of the sphenoid (Fig. 3C,I).

The detailed CNR results for differentiation of white and gray matter in the brain are listed in Suppl. Table 2. Using HiP-CT at 16.5 µm voxel size, the CNR for this differentiation is with 3.03 about 50 % lower than in MRI (CNR 6.03) due to the increased noise level that comes with higher resolution. However, when binning the data to a similar voxel size as the MR scan (HiP-CT: 265×265×265 µm^3^; MR: 214×214×225 µm^3^), the CNR for HiP-CT increased to 8.58, which is about 40 % higher than the CNR for MR.

### Structure tensor analysis of the brain

Figure 4 depicts the results of the structure tensor analysis of the brain. The vector field, re-oriented with respect to the anterior commissure – posterior commissure landmarks reveals similar structures to the human neuroanatomy, such as the cortico-spinal tract, the corpus callosum and the striatal fiber bundles.

**Figure 4:**
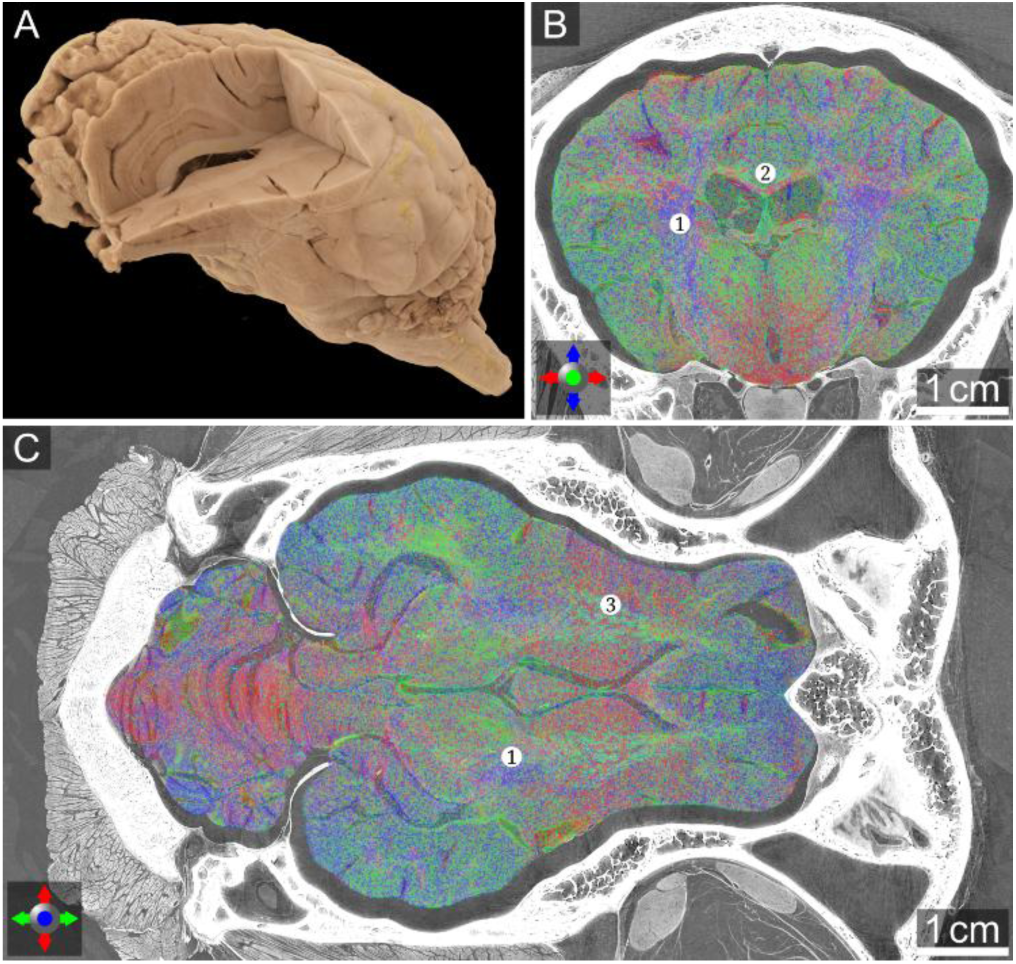
Structure tensor analysis of the brain. A, Segmented brain rendered in Cinematic Anatomy. B, C, Primary vector directions in the brain, along the coronal plane (B) and along the transverse plane (C), showcasing two major white matter bundles: the cortico-spinal tract in blue ❶ and the corpus callosum in red ❷, and smaller parallel bundles in green ❸ within the striatal region.

### Visualization of fine structures in the eye

Figure 5 shows the local tomography zoom of the left eye with 4.23 µm voxel size. HiP-CT with ethanol preparation demonstrates a high level of anatomical detail. Most notably, the ciliary body is visualised with exceptional clarity, displaying its fine structural differentiation and vascular patterns (Fig. 5B,D). The choroid and its microvascular architecture are clearly visible (Fig. 5E). The retinal layers, already well-studied by optical coherence tomography, are revealed with enhanced contrast and delineation (Fig. 5E). Furthermore, the optic nerve head, its microvasculature and its surrounding structures are rendered with high fidelity (Fig 5C,F), offering new insights into axonal and peripapillary morphology.

**Figure 5:**
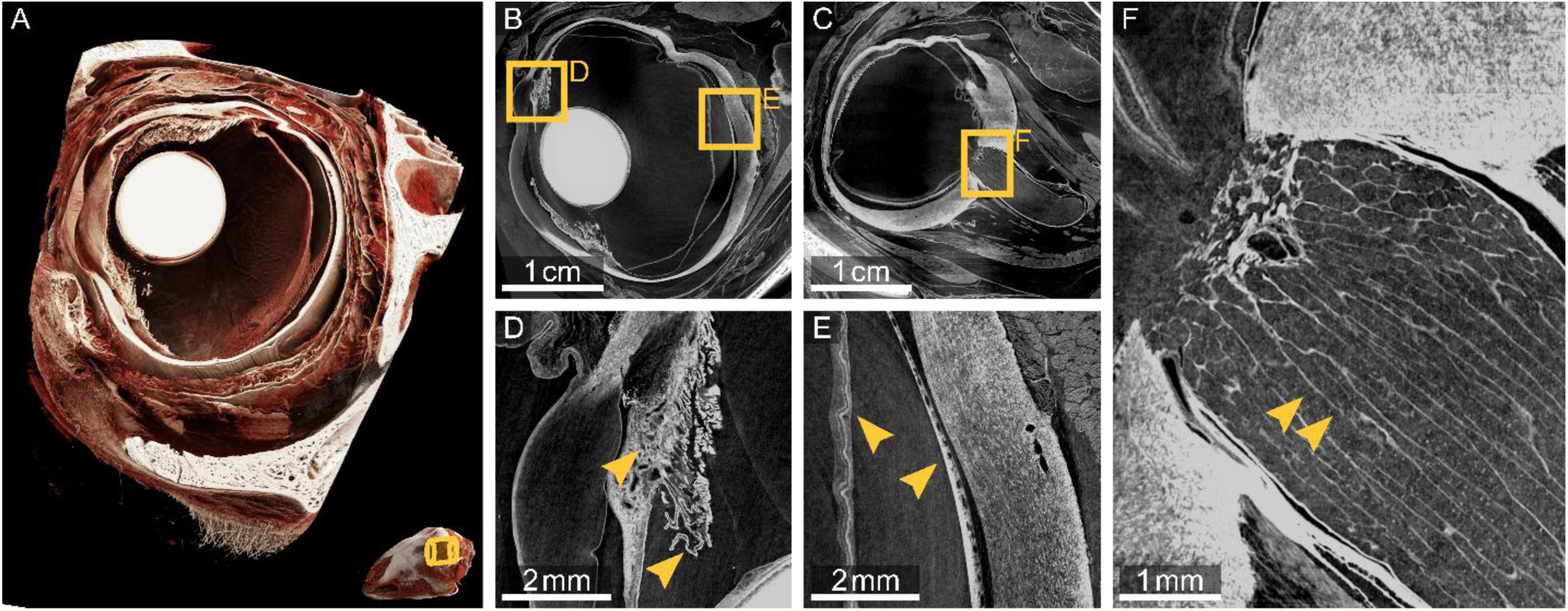
Local tomography in the eye with a voxel size of 4.23 μm. A, Volume rendered in Cinematic Anatomy with a clipping plane through the eye. B-F, exemplary slices with detail views to the regions indicated by the rectangles. Features of few microns like the layers of the retina, vasculature in the choroid and sclera and the bundles in the optic nerve can be distinguished.

### Structural and fibre organization of the coronal suture

Figure 6 shows the local tomography of the coronal suture with a voxel size of 2.20 µm and the associated collagen fibre bundle analysis. The median suture thickness of the coronal suture is 162 µm, with increased thickness around the edges of the suture at both dorsal and ventral suture edges.

**Figure 6:**
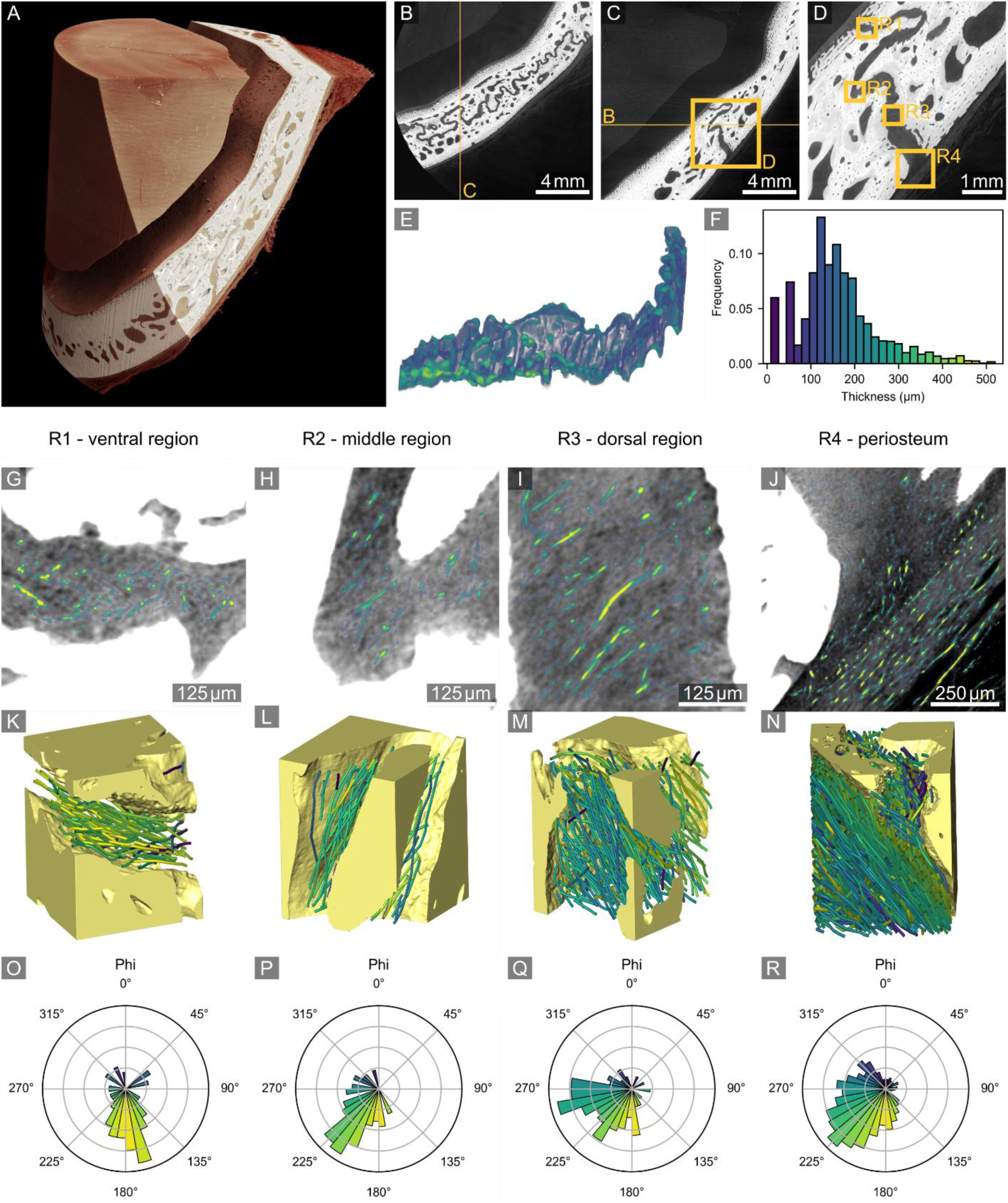
Local tomography in the coronal suture with a voxel size of 2.20 µm. A, Volume rendered in Cinematic Anatomy with a clipping plane to visualize the suture. B-D, exemplary slices. The rectangles R1 – R4 mark the subvolumes evaluated for collagen fibre bundle orientation. E, segmented suture color-coded by thickness. F, thickness distribution in the suture. G-J, cylinder correlation maps of the four regions. K-N, fibre bundle visualization with different rotations around the z-axis (vertical) for better visibility. O-R, angular distribution of fibre bundle orientation within the x-y plane (horizontal in K-N).

The periosteum is known to show dense, uniformly oriented fibres^23^. This organisation can be qualitatively observed in the periosteum analysis (Fig. 6J,N,R), Note that all artefacts from bones are streaks in the *x*-*y* plane, due to rotation around the *z*-axis during acquisition. Thus, if the fibre bundle analysis was tracking artefacts, these artefactual bundles would have a theta angle of 90° (compare Suppl. Fig. S2).

Suture fibre bundle orientation analyses reveals differences between the three investigated suture regions. Specifically, fibre bundles in the middle region show the highest homogeneity in orientation. Qualitatively, the fibre bundles largely run along the suture in parallel to the bone as opposed to clearly joining the two bones together. As the region immediately next to the bone was not considered in these analyses, it could not be analysed if or where the fibre bundle ends connect to the bone. Note that this sheep was a young animal, as it still has lacteal anterior teeth and developing third molars. Thus, it is to be expected that the cranial suture is not yet very stiff.

## Discussion

In this work we apply HiP-CT in combination with EPR to image a complete sheep head with a voxel size down to 16.5 µm and high contrast in both soft tissue and bone. Additionally, we show local tomography zooms with voxel sizes down to 2.20 µm in regions of interest. These zoom scans allow detailed analysis of structures of few microns, such as the layers of the retina and the collagen organisation within the cranial sutures.

This proof-of-principle HiP-CT scan of an entire sheep head prepared in 70% ethanol demonstrates the technical feasibility of capturing the intricate three-dimensional micro-anatomy of both soft and hard tissues in a non-destructive way.

HiP-CT imaging of excised soft-tissue organs such as the brain^5^ or hard-tissue samples such as the cochlea^24^ has been performed before. Due to less absorbing tissue, less dose is necessary and image quality in excised and smaller samples with either hard or soft tissue, or both, is usually better than for heterogeneous and bigger samples. However, for a complete understanding of the morphology it is often necessary to image the full sample without destroying it.

A whole human head has previously been imaged for dentistry applications with attenuation-based µCT with a voxel size of 140 µm and with very limited soft tissue contrast^25^. While soft tissue contrast in attenuation-based µCT can be improved with contrast agents, this increases complexity in sample preparation. In case of large ex-vivo samples, uniform contrast agent distribution throughout the sample is difficult to achieve. Using phase-contrast imaging increases soft-tissue contrast but poses high demands on the X-ray setup. Currently, laboratory-based techniques for phase-contrast imaging are not yet suitable for samples as large and dense as a head, mainly due to limitations in beam energy and sensitivity. Instead, it is necessary to resort to using synchrotron facilities. Nevertheless, using phase-contrast imaging for larger biomedical samples with laboratory x-ray sources is an active field of research^26–29^.

The soft-tissue contrast with HiP-CT combined with ethanol preparation allows differentiation of tissues with very similar attenuation, such as grey and white matter in the brain. The CNR analysis shows the high soft-tissue contrast even in a brain enclosed by the skull, and at much higher resolution than in clinical imaging. However, it has to be noted that the advantages of HiP-CT over clinical imaging come with the sample preparation, a high dose preventative of in-vivo imaging, and the high cost of a synchrotron x-ray source.

The zoom scans in the eye and the coronal suture showcase the possibility of distinguishing few-micron scale features in an intact sample. The ex-vivo detailed analysis of collagen fibre orientation and suture morphology at a few micrometers resolution provides novel insights into cranial suture architecture, which may improve surgical planning, graft integration, and diagnosis in craniofacial surgery, trauma reconstruction, and conditions involving abnormal suture development such as craniosynostosis.

HiP-CT bridges a critical gap in ex-vivo anatomical imaging: it combines cm-scale context with microscopic detail. This work advances the achievable field of view and resolution on heterogeneous biomedical samples, thus broadening the possible applications of HiP-CT. It paves the way for applications in other anatomically complex regions where the interplay between soft and bony tissues is essential, such as the pelvis or even an entire fetus, allowing unprecedented investigation of developmental anatomy, congenital anomalies, and musculoskeletal pathology.

Although further technical developments are required, this is a significant step towards applying HiP-CT to the larger human head. The larger sample size will entail a higher X-ray dose and thus a higher complexity in sample preparation and required stability. If successful, such advancement would offer significant clinical implications for radiology, ear-nose-throat, ophthalmology, neurosurgery, maxillofacial surgery, dentistry and all other medical specialties concerned with head and cranial pathology.

There are limitations to using HiP-CT for studying heterogeneous biomedical samples. A high dose level is required to obtain a high contrast with such small voxel sizes. Especially for the brain, the preparation with ethanol is required to improve contrast, and preparation times may be long for large samples to achieve degassing and ethanol equilibrium. Additionally, the preparation might induce changes in the sample’s morphology, e.g. dehydration with ethanol may make the soft tissue shrink. Access to a synchrotron facility and the sample preparation limit throughput and scalability. And finally, the data sizes of HiP-CT can be easily in the Terabyte range for a single volume, making the data visualization and analysis challenging. These factors make it hard to conduct studies with larger sample cohorts for statistical significance of the results.

In conclusion, HiP-CT with ethanol preparation and EPR allows non-destructive, multiscale imaging of a complete sheep head with microstructural detail. This enables its morphological analysis from full anatomy to few-micron scale features, opening the possibility for further investigations on hard-soft tissue interfaces and on soft tissues embedded in hard tissues.

## Abbreviations

HiP-CT: Hierarchical Phase-Contrast Tomography
MRI: Magnetic Resonance Imaging
CT: X-ray Computed Tomography
µCT: Micro-Computed Tomography
CNR: Contrast-to-noise ratio

## Acknowledgements

We gratefully acknowledge ESRF-The European Synchrotron beamtime md1290 on BM18 as the data source and the BM18 team for the support. We thank the Radiology Department of the University Hospital of Grenoble (CHUGA) for their time and effort to perform the clinical imaging. We thank David Stansby and Guillaume Gaisne for their support with data management and uploading.

## Supplementary Material

### Movie captions

#### Movie 1: Cinematic whole head overview

Cinematic 3D rendering of the overview scan (16.5 µm voxels) of the complete head, with changing virtual clipping planes to highlight structures within the head such as the brain, the tongue, and the eyes. Movie created by Theresa Urban using Cinematic Anatomy (Siemens Healthineers, Germany).

#### Movie 2: Comparison of HiP-CT and MR

Aligned rendering of HiP-CT overview scan (27.7 µm voxels) and clinical MR images. HiP-CT provides soft-tissue contrast similar to that of MR, but with much higher spatial resolution. Movie created by Paul Tafforeau using VGSTUDIO MAX (Volume Graphics GmbH, Germany).

### Data availability and neuroglancer links

All volumes mentioned is this manuscript are available for download in BioImageArchive. This archive is currently accessible only via the provided link for review, but will be made public upon acceptance.

Neuroglancer enables visualization of data through a web browser, without the need to download to a local disk. Neuroglancer links to all HiP-CT volumes are available in the provided csv file. See https://neuroglancer-docs.web.app/user-guide/navigation.html for instructions on how to use neuroglancer.

### HiP-CT acquisition parameters

**Supplementary Table 1:** All HiP-CT volumes with their context and acquisition parameters. Abbreviations: refl., reflective; Proj., Projections; Transl., Translation; Prop., Propagation; px, pixel. See metadata of volumes on BioImageArchive for more detailed parameters.

| Context | Voxel size | Scintillator | Optic, magn. | Camera | Detector ROI | Lateral FOV | Prop. Distance | Mode | Proj. per turn | Transl. per turn | Attenuators | Mean Energy | Imaging time | Imaging date |
| --- | --- | --- | --- | --- | --- | --- | --- | --- | --- | --- | --- | --- | --- | --- |
| Overview | 27.7 $\mu\text{m}$ | LuAg<br>2000 $\mu\text{m}$ | 0.175x | IRIS 15 | 3104 px ×<br>256 px | 147 mm | 30 m | zseries | 8000 | 5 mm | 0.44 mm Mo | 114 keV | 4.1 h | Nov 20<sup>th</sup> 2022 |
| Overview | 16.5 $\mu\text{m}$ | LuAg<br>2000 $\mu\text{m}$<br>refl. layer | 0.125x | IRIS 15 | 5056 px ×<br>600 px | 149 mm | 29.4 m | helical | 15000 | 5 mm | 10 mm Al<sub>2</sub>O<sub>3</sub><br>0.3 mm Ag | 117 keV | 1.7 h | Jun 21<sup>th</sup> 2024 |
| Cochlea | 2.20 $\mu\text{m}$ | LuAg<br>250 $\mu\text{m}$<br>refl. layer | 2x | IRIS 15 | 5056 px ×<br>2960 px | 20 mm | 2.5 m | zseries | 15000 | 5.5 mm | 1.3 mm Mo | 119 keV | 1.4 h | Dec 7<sup>th</sup> 2023 |
| Upper molars | 2.20 $\mu\text{m}$ | LuAg<br>250 $\mu\text{m}$<br>refl. layer | 2x | IRIS 15 | 5056 px ×<br>2960 px | 20 mm | 2.5 m | zseries | 15000 | 5.5 mm | 1.3 mm Mo | 119 keV | 2.1 h | Dec 7<sup>th</sup> 2023 |
| Lower molars | 2.20 $\mu\text{m}$ | LuAg<br>250 $\mu\text{m}$<br>refl. layer | 2x | IRIS 15 | 5056 px ×<br>2960 px | 20 mm | 2.5 m | zseries | 15000 | 5.5 mm | 1.3 mm Mo | 119 keV | 2.6 h | Dec 8<sup>th</sup> 2023 |
| Nose turbinate | 2.20 $\mu\text{m}$ | LuAg<br>250 $\mu\text{m}$<br>refl. layer | 2x | IRIS 15 | 5056 px ×<br>2960 px | 20 mm | 2.5 m | zseries | 15000 | 5.5 mm | 1.3 mm Mo | 119 keV | 2.1 h | Dec 8<sup>th</sup> 2023 |
| Incisors | 2.20 $\mu\text{m}$ | LuAg<br>250 $\mu\text{m}$<br>refl. layer | 2x | IRIS 15 | 5056 px ×<br>2960 px | 20 mm | 2.5 m | zseries | 15000 | 5.5 mm | 1.3 mm Mo | 119 keV | 1.1 h | Dec 8<sup>th</sup> 2023 |
| Tongue | 2.20 $\mu\text{m}$ | LuAg<br>250 $\mu\text{m}$<br>refl. layer | 2x | IRIS 15 | 5056 px ×<br>2960 px | 20 mm | 2.5 m | zseries | 15000 | 5.5 mm | 1.3 mm Mo | 119 keV | 1.1 h | Dec 8<sup>th</sup> 2023 |
| Left eye | 4.23 $\mu\text{m}$ | LuAg<br>100 $\mu\text{m}$<br>refl layer | 1x | IRIS 15 | 5056 px ×<br>1400 px | 38 mm | 4.0 m | helical | 15000 | 3 mm | 10 mm Al<sub>2</sub>O<sub>3</sub><br>0.3 mm Ag<br>15 mm C | 100 keV | 1.3 h | Jun 21<sup>th</sup> 2024 |
| Coronal suture | 2.20 $\mu\text{m}$ | LuAg<br>250 $\mu\text{m}$<br>refl. layer | 2x | IRIS 15 | 5056 px ×<br>2960 px | 20 mm | 1.5 m | helical | 15000 | 3 mm | 1 mm Al<sub>2</sub>O<sub>3</sub><br>0.44 mm Mo | 99 keV | 1.2 h | Sept 18<sup>th</sup> 2024 |
| Optic chiasm | 2.20 $\mu\text{m}$ | LuAg<br>250 $\mu\text{m}$<br>refl. layer | 2x | IRIS 15 | 5056 px ×<br>2960 px | 20 mm | 1.5 m | helical | 15000 | 3 mm | 1 mm Al<sub>2</sub>O<sub>3</sub><br>0.44 mm Mo | 99 keV | 1.2 h | Sept 18<sup>th</sup> 2024 |
| Interfrontal suture | 2.20 $\mu\text{m}$ | LuAg<br>250 $\mu\text{m}$<br>refl. layer | 2x | IRIS 15 | 5056 px ×<br>2960 px | 20 mm | 1.5 m | helical | 15000 | 3 mm | 1 mm Al<sub>2</sub>O<sub>3</sub><br>0.44 mm Mo | 99 keV | 1.0 h | Sept 18<sup>th</sup> 2024 |
| Lambdoid suture | 2.20 $\mu\text{m}$ | LuAg<br>250 $\mu\text{m}$<br>refl. layer | 2x | IRIS 15 | 5056 px ×<br>2960 px | 20 mm | 1.5 m | helical | 15000 | 3 mm | 1 mm Al<sub>2</sub>O<sub>3</sub><br>0.44 mm Mo | 99 keV | 1.0 h | Sept 18<sup>th</sup> 2024 |

### CNR analysis

Regions in white matter, grey matter, and in the background ethanol were segmented using VGSTUDIO MAX (Volume Graphics GmbH, Germany). The agar in the support was excluded from the ethanol background segmentation, as it shows some structural contrast in HiP-CT. The contrast-to-noise ratio (CNR) was calculated as

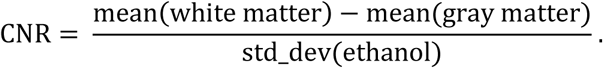

**Supplementary Table 2:**
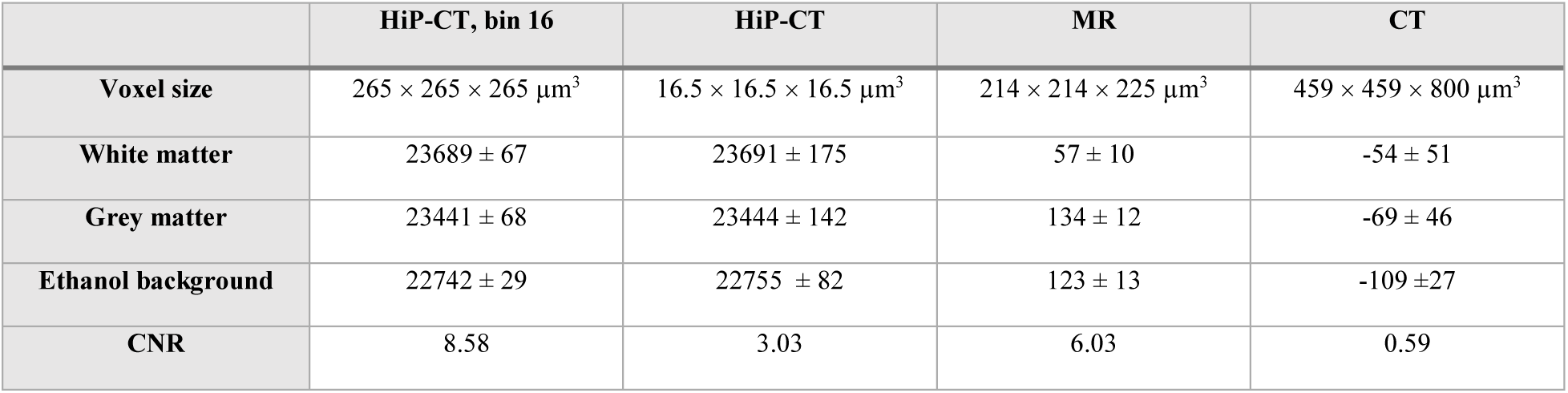
Signal values for CNR calculation. For HiP-CT, the CNR was calculated for both the full dataset and the dataset binned by 16, yielding a voxel size in the range of those of the clinical modalities. Values are given as mean ± standard deviation.

**Supplementary Figure S1:**
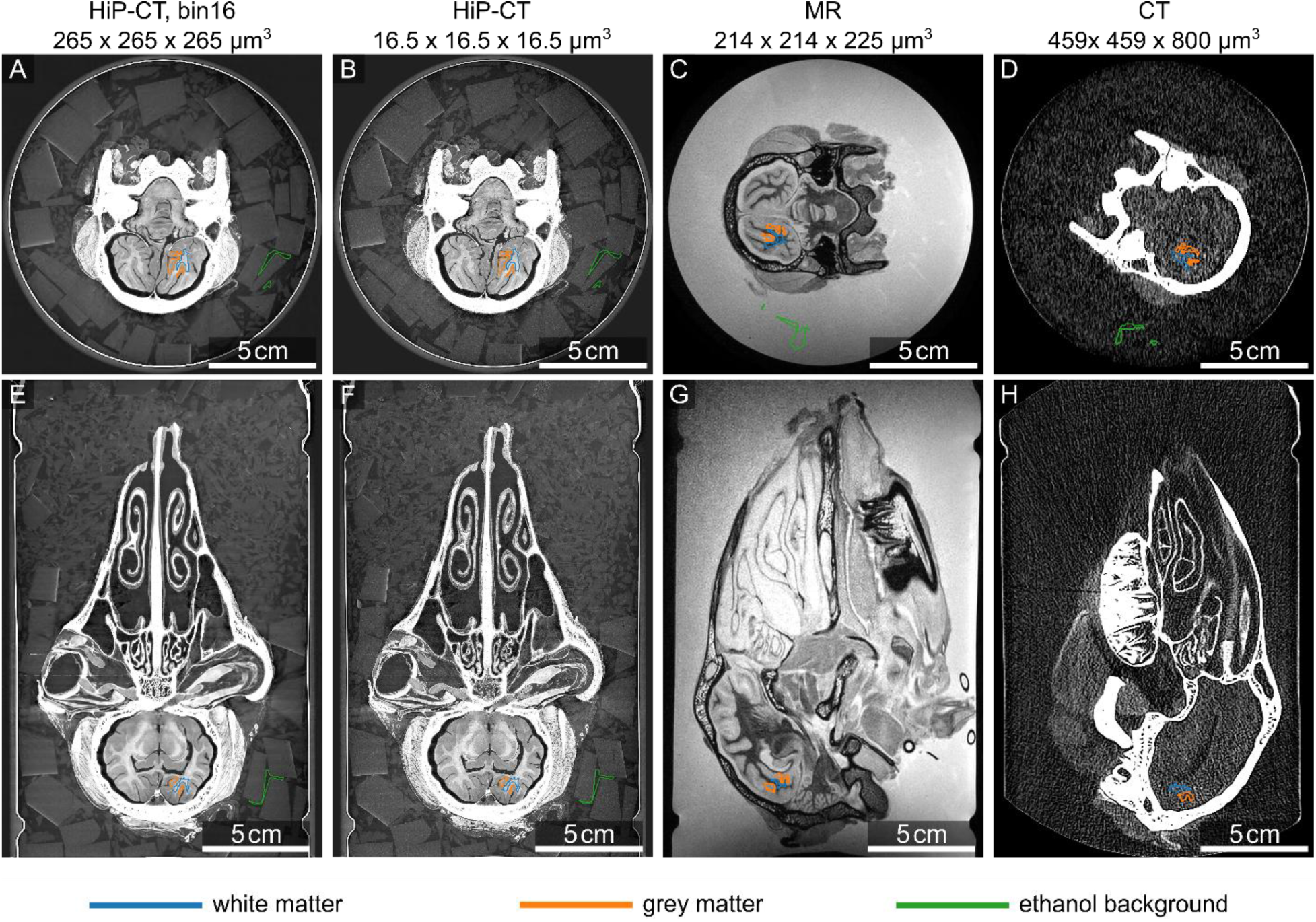
Segmentations for CNR calculation. The regions were segmented in the binned HiP-CT volume and transferred to the other modalities with the transforms from registration.

### Coronal Suture Segmentation

The BounTI algorithm^21^ was used for suture segmentation: Two disconnected segments for the frontal and parietal bones were produced via thresholding. Each bone segment was grown (expanded by one voxel at a time) until both segments contacted in the middle of the suture. Contact voxels were selected and grown until the new segment contacted the original bone; the new segment was then considered the suture. The segment was then manually edited at the suture/periosteum boundary, keeping the suture flush with the bones.

### Fiber Analysis

To enhance the contrast of collagen structures, 3D unsharp masking (edge size – 7, edge contrast – 0.7, brightness threshold – 0) was applied. Cylinder correlation^30–32^ was performed with the following parameters: Cylinder length 87 µm, angular sampling 10°, mask cylinder radius 12 µm, outer cylinder radius 7 µm and inner cylinder radius 0 µm. The correlation field and orientation fields were masked 10 voxels from each side, including the bone, to remove artefactual results that occur at the image boundaries and bone/suture interface.

Trace correlation line module was then used to obtain individual fibres with the following parameters: Minimum seed correlation 90, minimum continuation correlation 60, direction coefficient 0.5, minimum distance 25 µm, minimum length 25 µm. The following search parameters were used: length 87 µm, angle 37°, minimum step size 10%. Additionally, multiple hypothesis tracing^33^ was enabled with search depth and width set to 3.

**Supplementary Figure S2:**
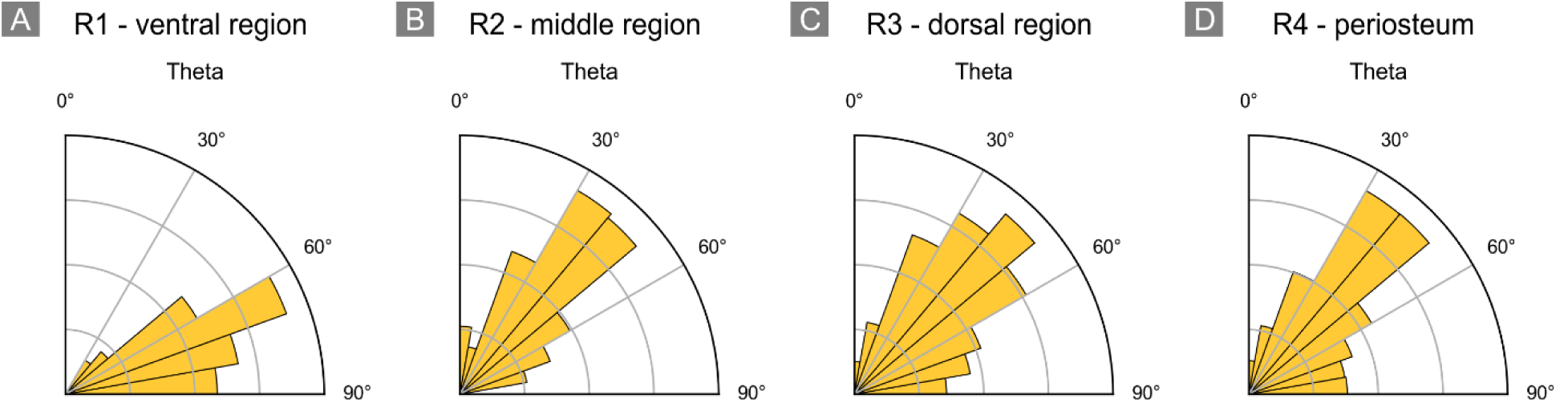
Theta angle distribution of fibre bundles in the four analyzed regions of the suture.

